# Information decomposition of multichannel EMG to map functional interactions in the distributed motor system

**DOI:** 10.1101/587949

**Authors:** Tjeerd W Boonstra, Luca Faes, Jennifer N Kerkman, Daniele Marinazzo

## Abstract

The central nervous system needs to coordinate multiple muscles during postural control. Functional coordination is established through the neural circuitry that interconnects different muscles. Here we used multivariate information decomposition of multichannel EMG acquired from 14 healthy participants during postural tasks to investigate the neural interactions between muscles. A set of information measures were estimated from an instantaneous linear regression model and a time-lagged VAR model fitted to the EMG envelopes of 36 muscles. We used network analysis to quantify the structure of functional interactions between muscles and compared them across experimental conditions. Conditional mutual information and transfer entropy revealed sparse networks dominated by local connections between muscles. We observed significant changes in muscle networks across postural tasks localized to the muscles involved in performing those tasks. Information decomposition revealed distinct patterns in task-related changes: unimanual and bimanual pointing were associated with reduced transfer to the pectoralis major muscles, but an increase in total information compared to no pointing, while postural instability resulted in increased information, information transfer and information storage in the abductor longus muscles compared to normal stability. These findings show robust patterns of directed interactions between muscles that are task-dependent and can be assessed from surface EMG recorded during static postural tasks. We discuss directed muscle networks in terms of the neural circuitry involved in generating muscle activity and suggest that task-related effects may reflect gain modulations of spinal reflex pathways.

## 1. Introduction

Adaptive behavior emerges in the nervous system from the ongoing interactions among the body and the environment. Coordinated adaptations in the nervous system and periphery are essential for motor function and are established by integrating feedforward commands from the nervous system and sensory feedback from the body (Chiel and Beer, 1997). It is argued that a balance between functional segregation and integration in the nervous system is required for perception and behavior (Tononi et al., 1994). These cognitive functions result from the dynamic interactions of distributed neural populations operating in large-scale networks that determine the flow of information through the central nervous system (Bressler and Menon, 2010; Sporns et al., 2004). While the functional implications of large-scale networks have mainly been investigated within the brain (Petersen and Sporns, 2015), we expected that similar principles apply to the entire nervous system and encompass interactions between brain and body.

The brain and spinal cord are interwoven with the body and interact through the peripheral and autonomic nervous systems with other organ systems (Freund et al., 2016). Through these neuronal pathways, dynamic interactions among subsystems are mediated to support physiological function and establish system-wide integration (Bashan et al., 2012). In the motor system, such functional integration involves the descending pathways that connect supraspinal motor areas to the spinal motor neurons generating the muscle activation patterns. The corticospinal tract originates from different cortical areas including primary motor cortex and terminates widely within the spinal gray matter, including direct monosynaptic projections to contralateral spinal motor neurons (Lemon, 2008). Other descending pathways such as the vestibulospinal and the reticulospinal tract originate from brainstem nuclei, project to bilateral regions of the spinal cord and are involved in controlling posture and locomotion (Kuypers, 1981). In addition to divergent projects from supraspinal regions, propriospinal pathways and spinal interneurons interconnect motor neuron pools innervating different muscles (Pierrot-Deseilligny and Burke, 2005). Finally, through the mechanical couplings within the musculoskeletal system, motor neurons receive afferent feedback from the activity of other muscles (Kutch and Valero-Cuevas, 2012). Through all these pathways, focal motor activity is expected to generate a cascade of muscle activations that need to be controlled to ensure coordinated movements. The dynamic interplay between sensory feedback and motor output is essential for motor coordination and gain modulation is an important mechanism to control the flow of activity through the sensorimotor system (Azim and Seki, 2019; Krakauer, 2019; Pierrot-Deseilligny and Burke, 2005; Pruszynski and Scott, 2012). However, gain modulation in sensorimotor pathways is difficult to assess during natural movements.

Information theory provides the foundation for investigating the processing and flow of information in distributed functional networks. In particular, information dynamics provide a versatile and unifying set of tools to dissect general information processing into basic elements of computation which quantify the new information produced in a network at each moment in time as well as the amounts of information stored and transferred within the network (Deco and Schürmann, 2012; Faes et al., 2017b; Lizier, 2012). Temporal precedence is the key concept that allows to quantify directed interactions between individual components of a large-scale system. For example, Granger causality is a statistical method to determine whether one time series can be used to predict another (Granger, 1969). It has been widely used in neuroscience research to characterize functional brain circuits (Seth et al., 2015). Granger causality is closely linked to transfer entropy, an information-theoretic measure of time-directed information transfer (Barnett et al., 2009). Despite their widespread application, there are ongoing discussions regarding the interpretation of these methods in neuroscience research (Barnett et al., 2018; Faes et al., 2017c; Stokes and Purdon, 2017). An issue with Granger causality measures is that they are often estimated using a subset of variables, though being likely influenced by other variables, either available or exogenous (unrecorded). This issue can be partially circumvented by estimating conditional information transfer (Guo et al., 2008; Runge et al., 2012) and by investigating how multiple source variables interact with each other while they transfer information towards a target in the network (Stramaglia et al., 2014). Moreover, it has been argued that multiple information-theoretic measures, derived from tools such as information decomposition (Faes et al., 2017b), should be combined to provide an exhaustive view of the underlying dynamics of brain and physiological networks.

In this paper, we use information decomposition to map interactions in functional muscle networks involved in postural control. The musculoskeletal system consists of more than 300 skeletal muscles, which are controlled by the central nervous system through the spinal motor neurons. The activity of motor neurons can be noninvasively recorded using surface electromyography (EMG) and intermuscular (EMG-EMG) coherence is then used to investigate functional connectivity in the sensorimotor system (Farmer, 1998). Intermuscular coherence is thought to reflect correlated inputs to spinal motor neurons that may have different origins (Boonstra et al., 2016). Using a formalism of information decomposition of target effects from multi-source interactions, we aim to disentangle the neural sources that contribute to correlated inputs to spinal motor neurons. We apply this approach to EMG activity recorded from multiple muscles distributed across the body while participants performed postural tasks. We previously used intermuscular coherence to map undirected functional muscle networks in this data (Kerkman et al., 2018). Here, we use conditional transfer entropy to map directed interactions between muscles and investigate their contribution in the coordination of the neuromuscular system during postural control.

## 2. Material and Methods

### 2.1 Experimental protocol and data acquisition

We used multi-channel EMG data that was acquired from fourteen healthy participants (seven males and seven females; mean age, 25 ± 8 years; ten right-handed and four left-handed) while standing upright (Kerkman et al., 2018). The experiment was approved by the Ethics Committee Human Movement Sciences of the Vrije Universiteit Amsterdam (reference ECB 2014-78) and participants gave informed consent before participation.

The experiment consisted of nine experimental conditions in which postural stability and pointing behavior were varied using a full-factorial design. Postural stability was manipulated using a balance board (normal standing, instability in anterior-posterior direction and instability in medial-lateral direction) and participants held either their arms alongside their body (no pointing) or pointed a laser pointer on a target using either their dominant hand or both their hands. Each task was repeated in six trials of 30 s each and the experiment lasted about 1.5 hour including rests.

Bipolar surface EMG was recorded using Porti systems (TMSi, the Netherlands) from 36 muscles distributed across the body, i.e. eighteen bilateral muscles (Table 1). Electrode locations were based on SENIAM guidelines. Muscle that were not available in the SENIAM guidelines were localized based on palpation. EMG signals were online high-pass filtered at 5 Hz and sampled at 2 kHz. EMG data were offline band-pass filtered (1 - 400 Hz) before independent component analysis was used for electrocardiography removal (Willigenburg et al., 2012). Additionally, EMG signals were high pass filtered (20 Hz) and EMG envelopes were extracted using the Hilbert transform (Boonstra and Breakspear, 2012).

**Table 1.** List of muscles

|  | **muscle** | **abbreviation** |
| --- | --- | --- |
| 1 | tibialis anterior | TA |
| 2 | gastrocnemius medialis | GM |
| 3 | soleus | SOL |
| 4 | rectus femoris | RF |
| 5 | biceps femoris | BF |
| 6 | vastus lateralis | VL |
| 7 | adductor longus | AL |
| 8 | external oblique | EO |
| 9 | pectoralis major | PMA |
| 10 | sternocleidomastoideus | SMA |
| 11 | longissimus | LO |
| 12 | latissimus dorsi | LD |
| 13 | trapezius (transverse part) | TZ |
| 14 | deltoid (acromial part) | D |
| 15 | biceps brachii | BB |
| 16 | triceps brachii | TRB |
| 17 | extensor digitorum | ED |
| 18 | flexor digitorum superficialis | FDS |

Prior to information-theoretic analysis, the signals were down-sampled to 200 Hz, obtaining time series of length N=6000 points. Moreover, to further the fulfillment of stationarity criteria, the time series were detrended using a zero-phase autoregressive high-pass filter with cutoff frequency of 1.56 Hz (3 dB) (Nollo et al., 2000), and were then reduced to zero mean.

### 2.2 Information-theoretic concepts

The entropy measures used in this study to characterize muscle networks are based on basic information-theoretic concepts (Cover and Thomas, 2012), which are briefly recalled in the following. The information content of a scalar (one-dimensional) continuous random variable *X*, with probability density function *f*_*X*_(*x*), is quantified by its entropy, 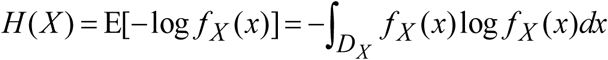, where E is the expectation operator, *D*_*X*_ is the domain of *X* and log is the natural logarithm so that entropy is measured in “nats”. If we consider a second *k*-dimensional vector variable *Z*=[*Z*_1_…*Z*_*k*_] with probability density *f*_*Z*_(*z*), the conditional information of *X* given *Z*, i.e. the information remaining in *X* when *Z* is known, is quantified by the conditional entropy *H* (*X*| *Z*) = E[− log *f*_*X*|*Z*_(*x* | *z*)], where *f*_*X*|*Z*_(*x*|*z*) denotes the conditional probability of *x* given *z*. The concepts of entropy and conditional entropy can be combined to measure the mutual information (MI) between *X* and *Z*, quantifying the information shared between the two variables, as *I*(*X*;*Z*)=*H*(*X*)–*H*(*X*|*Z*). Moreover, the conditional mutual information between *X* and *Z* given a third variable *U*, *I*(*X*;*Z*|*U*), quantifies the information shared between *X* and *Z* which is not shared with *U*, defined as *I*(*X*;*Z*|*U*)=*H*(*X*|*U*)−*H*(*X*|*Z*,*U*).

A viable approach to quantify these information measures is based on linear regression models. It is indeed known that, when the random variables have a joint Gaussian distribution, entropy and conditional entropy can be formulated analytically as:

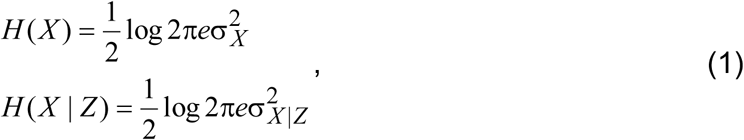

where 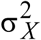 is the variance of the variable *X* and 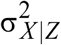 is the so-called partial variance, i.e. the variance of the residuals obtained performing a linear regression of *X* on all components of the vector variable *Z* (Barrett et al., 2010). Note that, since mutual information and conditional mutual information can be expressed as the sum of entropy and conditional entropy terms, Eq. (1) can be exploited to derive an analytical formulation for all information measures defined above.

### 2.3 Network information measures

The above information-theoretic preliminaries were exploited in this study to devise a signal processing framework for the description of how information is processed by muscle networks (Faes et al., 2017b). In such a framework, the EMG signals acquired from different muscles across the human body were interpreted as realizations of an *M*-dimensional vector stochastic process *Y*_*n*_=[*Y*_1,*n*_…*Y*_*M*,*n*_]^T^, where *n* denotes the current time sample (*M*=36 in this study, Fig. 1). The total information contained in the generic process *Y*_*j*_ at time *n*, as well as the information shared instantaneously (with zero lag) between two processes *Y*_*j*_ and *Y*_*i*_(*i*,*j*∈{1,…,*M*}) and the conditional information shared instantaneously by *Y*_*j*_ and *Y*_*i*_ given the other processes *Y*_*k*_ (*k*=[{1,…,*M*}/{*i*,*j*}]), are defined by the entropy, mutual information and conditional mutual information measures defined as:

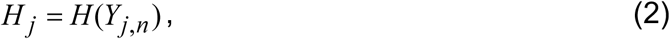

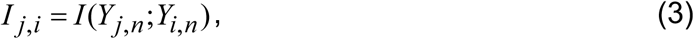

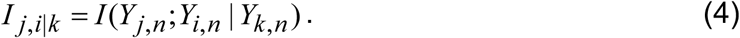

**Figure 1.**
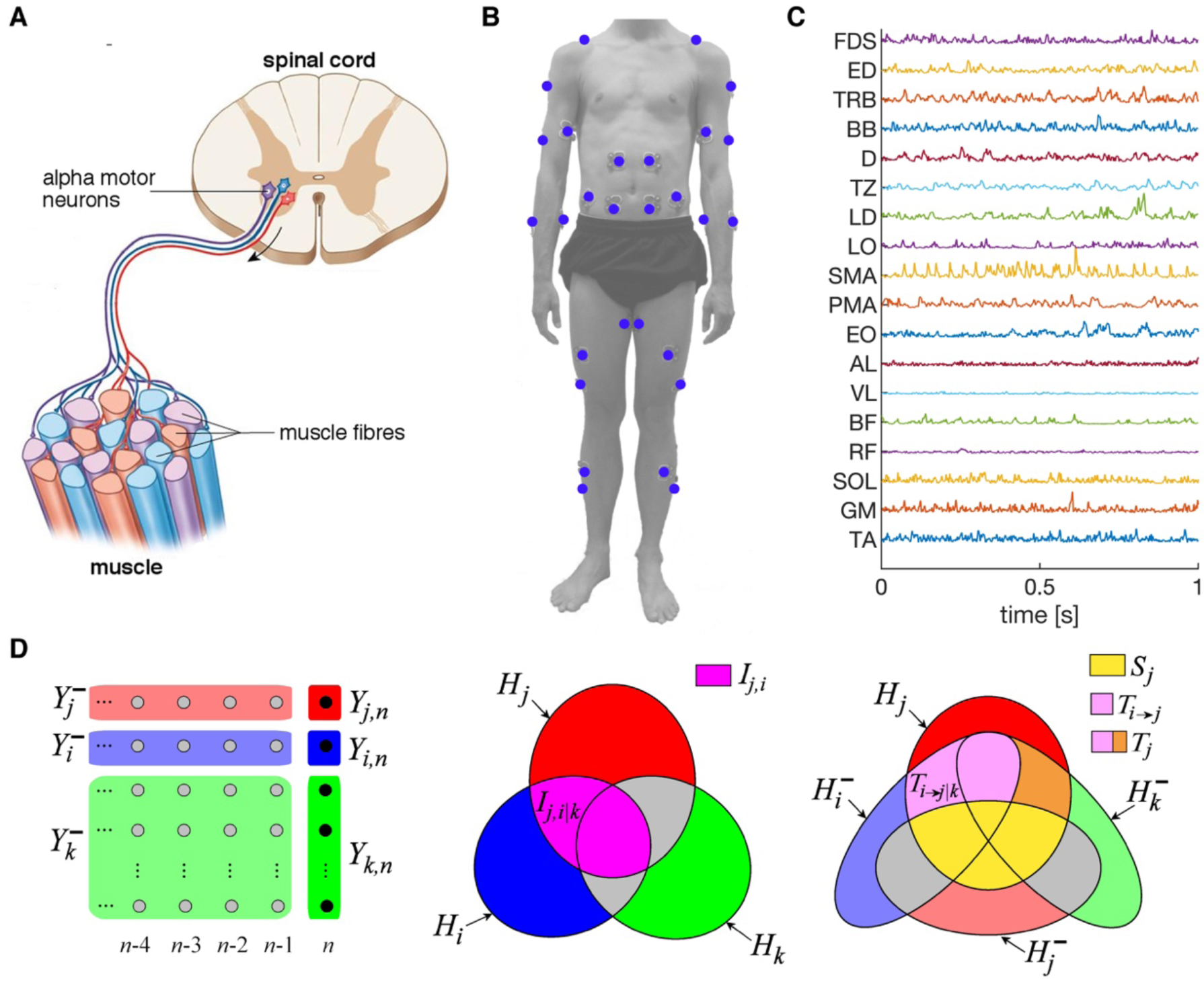
Overview of data acquisition and analysis steps. **A)** Motor units consist of alpha motor neurons located in the spinal cord that innervate muscle fibers. While EMG is recorded from the skin over the muscle, the EMG signals reflect neuronal activity generated in the spinal cord. **B)** Surface EMG is recorded from 36 muscles distributed across the body while participants performed postural control tasks. **C)** EMG data from eighteen muscles on the dominant side of the body during the bimanual pointing task from a single participant. **D)** Variables used to compute information measures: present (red) and past values (light red) of the target; present (blue) and past values (light blue) of the source; present (green) and past values (light green) of the remaining processes. The Venn diagram in the middle depicts instantaneous information metrics: mutual information between the target and the source is depicted in pink and the labeled part gives the conditional mutual information through which common effects from other processes are conditioned out. The Venn diagram on the right depicts time-lagged information metrics: information storage is depicted in yellow, transfer entropy is depicted in pink, conditional transfer entropy is the labeled part of the pink area, and total transfer entropy is the sum of the pink and orange areas.

A Venn diagram representation of how the above measures are obtained in terms of shared information is reported in the middle panel of Figure 1D. Moreover, defining as 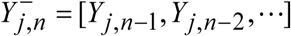 the (infinite dimensional) vector variable which represents the past history of the process *Y*_*j*_, the information stored in the *j*-th process is defined as the mutual information between its present and past variables:

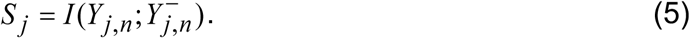

Consideration of past histories (Fig. 1D) also allows to quantify dynamic (time-lagged) causal interactions occurring from a source process *Y*_*i*_ toward a target process *Y*_*j*_. Specifically, the total information transferred to the target from all other processes in the network is computed as the conditional mutual information between the present of *Y*_*j*_ and the past of *Y*_*i*_ and of *Y*_*k*_, given the past of *Y*_*j*_:

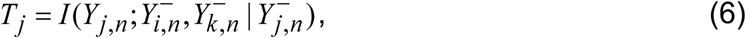

Then, with specific reference to one source process *Y*_*i*_, the bivariate information transfer from the source to the target, denoted as transfer entropy (TE), is measured as the conditional mutual information between the present of *Y*_*j*_ and the past of *Y*_*i*_ given the past of *Y*_*j*_:

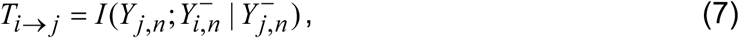

while the conditional information transfer is measured by conditioning also on the past of all other processes in the network collected in *Y*_*k*_:

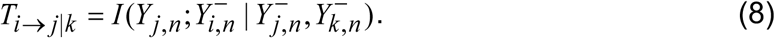

Note that, while each information measure is formulated accounting for the current time instant *n* in the right-hand side of its defining Eqs. (2–8), under the assumption of stationarity the measure is time-invariant so that the time index *n* can be omitted in the definition given in the left-hand side.

The Venn diagram in the right panel of Figure 1D shows a graphical representation of how the information shared by the present of the target and the past of the various processes in the motor system is combined to obtain the information storage as well as the total, bivariate and conditional information transfer.

### 2.4 Estimation of information measures

In this study, we computed information measures within a model-based linear processing frame, according to the block diagram depicted in Figure 2. Within this frame, all measures defined in Eqs. (2–8) were first expressed as sums of entropy and conditional entropy terms according to the definitions given in Sect. 2.2, and then computed using the formulations given in Eq. (1). To estimate entropy, the variance of each scalar process *Y*_*j*_ associated with the *j*-th EMG channel, 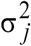, was estimated from the corresponding EMG recording. To estimate conditional entropy, the partial variances to be used in Eq. (1) were estimated from linear regression models.

**Figure 2.**
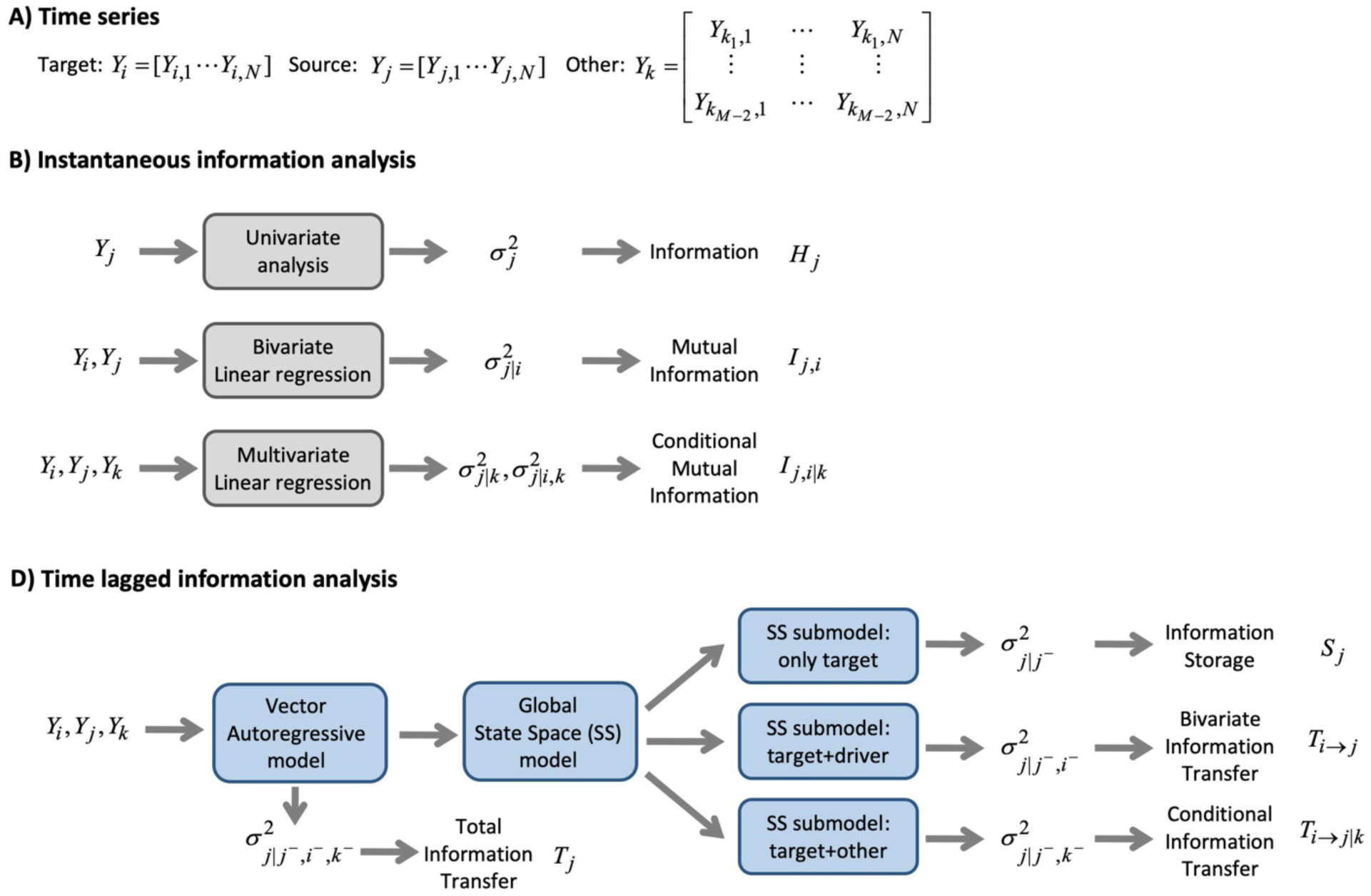
Block diagrams of the computational steps for estimating information measures. **A)** The target and source processes in a set of *M* time series (EMG signals) each composed of *N* observations. **B)** Estimation of instantaneous information measures. **C)** Estimation of time-lagged information measures.

Specifically, starting from a set of *M* time series (EMG signals) each composed of *N* observations (Fig. 2A), instantaneous information measures were computed as depicted in Figure 2B. The information content of the target time series *Y*_*j*_, was obtained simply as a function of its variance 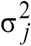.To estimate instantaneous mutual information, the partial variance of *Y*_*j*_ given *Y*_*i*_, 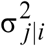, was estimated as the variance of the prediction error obtained regressing the *j*-th EMG channel on the *i*-th channel, and was used in conjunction with the variance of the *j*-th channel 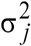 to compute the mutual information of Eq. (3). To estimate instantaneous conditional mutual information, the partial variances of *Y*_*j*_ given *Y*_*k*_,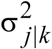, and of *Y*_*j*_ given both *Y*_*i*_ and *Y*_*k*_, 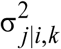, were estimated as the variance of the prediction errors obtained regressing the *j*-th EMG signals respectively on all other signals except the *i*-th, and on all other signals including the *i*-th, and used to compute the conditional mutual information of Eq. (4).

The time-lagged measures of Eqs. (2–8) were computed implementing the concept of Granger causality in a state space formulation (Barnett and Seth, 2015; Faes et al., 2017a) following the steps depicted in Figure 2C. Specifically, first linear regression was performed jointly for all processes to identify a vector autoregressive (VAR) model from the available multivariate EMG signals. The VAR model order, corresponding to the number of samples used to cover the past of the processes, was set according to the Bayesian Information Criterion (BIC) (Schwarz, 1978). The variance of the prediction error, associated to the VAR description of the *j*-th EMG signal, was taken as an estimate of the partial variance of *Y*_*j*_ given the past of all processes, 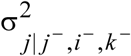. This estimated partial variance was used in conjunction with the variance of the target time series to estimate the total information transferred towards the target. We then used a state-space modeling approach (Barnett and Seth, 2015) to compute the remaining partial variances necessary to estimate the time-lagged information measures, i.e. the partial variance of *Y*_*j*_ given its own past, 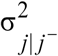, given its past and the past of *Y*_*i*_, 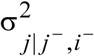,and given its past and the past of *Y*_*k*_, 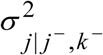. This method converts the VAR model into a state-space model and then rearranges the parameters such that submodels can be formed which contain only some of the processes. The partial variances relevant to these submodels can be computed analytically. With this approach it is possible to obtain computationally reliable estimates of all partial variances, which can then be used to compute the measures of information storage, and bivariate and conditional information transfer. We refer to Faes et al. (2017a) for the mathematical details about the state-space computation of the measures of information dynamics.

### 2.5 Statistical analysis

We mirrored the data of the left-handers to create a dominant and nondominant side of the body. We used network analysis to assess patterns of functional interactions between muscles and compared information measures across conditions. Mutual information and transfer entropy were estimated between all muscle pairs and provided the edge weights of the muscle networks. We used the weighted degree and in-degree to quantify the information that is shared with each target. We hence obtained 36 network metrics for each condition, trial and participant.

For statistical comparison across experimental conditions, we averaged the network metrics across the six trials per condition. In three participants, one trial was discarded due to bad electrode contact and for those participants network metrics were averaged across the remaining five trials. We then used a two-way ANOVA, with pointing behavior (no pointing, pointing with dominant hand, and pointing with both hands) and postural stability (normal standing, anterior-posterior instability, and medial-lateral instability) as independent factors, to statistically compare conditional transfer entropy across experimental conditions. We used Bonferroni-Holm correction to control the family-wise error rate. Alpha was set at 0.05.

We binarized the muscle networks for visualization purposes. The presence of binary edges was determined based on the statistical significance. As each of these measures was obtained comparing two nested linear regression models for any pair of channels, the statistical significance of the measure was tested using the Fisher F test comparing the residual sum of squares (Chow test; Chow, 1960). For every participant, we considered an edge as significant if the measure was significant in at least half of the trials (three of the five or six trials). At group-level, edges were considered significant if present in at least half of the participants (seven of the fourteen participants). The binary networks were then displayed on a model of the human body (Makarov et al., 2015) and the F-statistic defined at each node of the network were depicted as color-coded values interpolated on the body surface mesh (Jacobson, 2018).

## 3. Results

We used information decomposition of EMG activity measured from multiple muscles during postural control tasks and assessed mutual information to investigate functional dependencies at zero lag and transfer entropy to map the directed interactions between muscles accounting for the temporal flow of information. Furthermore, we estimated the conditional information metrics to account for common drive or cascade effects from other processes in the system. By comparing these information metrics across conditions, we investigated how these measures can be used to map changes in functional interactions in the sensorimotor system.

### 3.1 Mutual information

We first estimated mutual information between all muscle pairs in the bimanual pointing condition during normal stability, resulting in an adjacency matrix for every participant that was thresholded based on statistical significance to obtain undirected binary networks. To check these results, we also computed the adjacency matrices based on the correlation coefficient estimated directly from the EMG envelopes. After a transformation (−0.5*log(1-CC^2^)), the grand-average adjacency matrices are very similar (Mantel test, rho = 0.67, p = 1.6 * 10^−7^). The adjacency matrix showed high weights along the diagonal indicating edging between neighboring muscles (Fig. 3A, top panel). The strongest mutual information was observed between the D and TRB in both arms. Indeed, after thresholding, a sparse disconnected network (density = 0.13) was obtained showing primary connectivity between upper arm muscles but also separate cliques for the leg muscles (Fig. 3A, bottom panel). The nodes within each clique were densely connected, which may reflect the confounding influence of other processes. We therefore assessed conditional mutual information, which estimates the mutual information between two variables given all other variables and hence provides a better estimate of the true association between muscle pairs. The adjacency matrix showed reduced values for the conditional mutual information and the corresponding binary network was sparser (density = 0.09). However, the network structure remained largely unchanged (Mantel test, rho = 0.79, p = 1.0 * 10^−7^).

**Figure 3.**
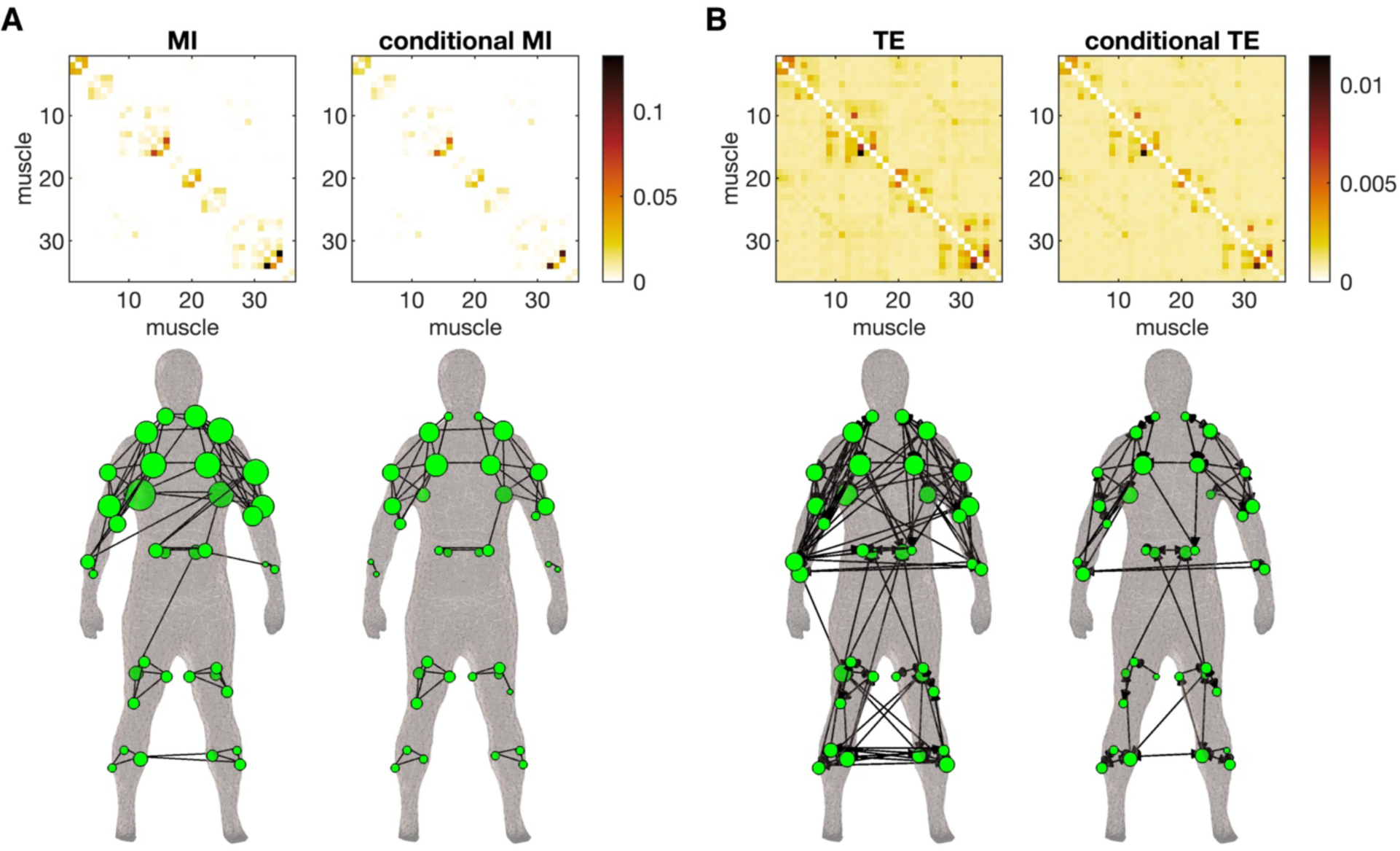
Undirected and directed muscle networks in bimanual pointing condition averaged across participants. **A)** Top panels show the grand-average adjacency matrices based on the mutual information (left) and the conditional mutual information (right). The order of the muscles in shown in Table 1 (first 16 muscles are on the right side and the last 16 muscles on the left). Bottom panels show the spatial topology of the binary networks displayed on the human body (Makarov et al., 2015). Lines represent significant edges between nodes (muscles), i.e. at least half of the participants showed significant mutual information. Node size represents the node degree. **B)** Top panels show the grand-average adjacency matrix based on the transfer entropy (left) and the conditional transfer entropy (right). Bottom panels show the spatial topology of binary networks. Arrows represent significant directed edges between nodes, i.e. at least half of the participants showed significant transfer entropy.

### 3.2 Transfer entropy

Next, we investigated information transfer to map directed interactions between muscles. Again, transfer entropy revealed mostly short-range connections as reflected by the high values of transfer entropy along the diagonal (Fig. 3B, top panel). After thresholding, a directed network was obtained showing mainly short-range connections but also some long-range connections between bilateral forearm muscles and between upper and lower body muscles (Fig. 3B, bottom panel; density = 0.14). We then assessed conditional transfer entropy to account for the influence of potential confounding influences. While the overall connectivity structure is similar to unconditioned transfer entropy, the network is much sparser (density = 0.05). Information transfer is mainly observed between upper arm and shoulder muscles, between bilateral leg and between lower arm muscles, but also a few edges interconnecting these cliques.

### 3.3 Experimental effects

We then compared the conditional transfer entropy across conditions. The directed networks showed a clear effect of the experimental manipulations. In the no pointing condition, only a few edges were observed between arm muscles (Fig. 4, left column), which is expected given the minimal EMG activity of those muscles in this condition. In contrast, when pointing with one arm, multiple edges could be observed in the arm used for pointing (the dominant right arm) but not in the contralateral arm (Fig. 4, middle column). In the bimanual pointing condition multiple edges could be observed in both arms (Fig. 4, right column). Similar effects could be observed between leg muscles in the postural conditions. In the normal standing condition, a few short-range connections were present between the leg muscles (Fig. 4, top row). When posture was destabilized in the anterior-posterior direction, the number of connections between leg muscles was greatly enhanced showing multiple edges between bilateral lower leg muscles and between upper and lower leg muscles (Fig. 4, middle row). Finally, when posture was destabilized in the medial-lateral direction, multiple connections were observed between the leg muscles, but now also many connections between leg muscles and muscles in the torso could be observed (Fig. 4, bottom row). We hence observed systematic changes in conditional transfer entropy across conditions.

**Figure 4.**
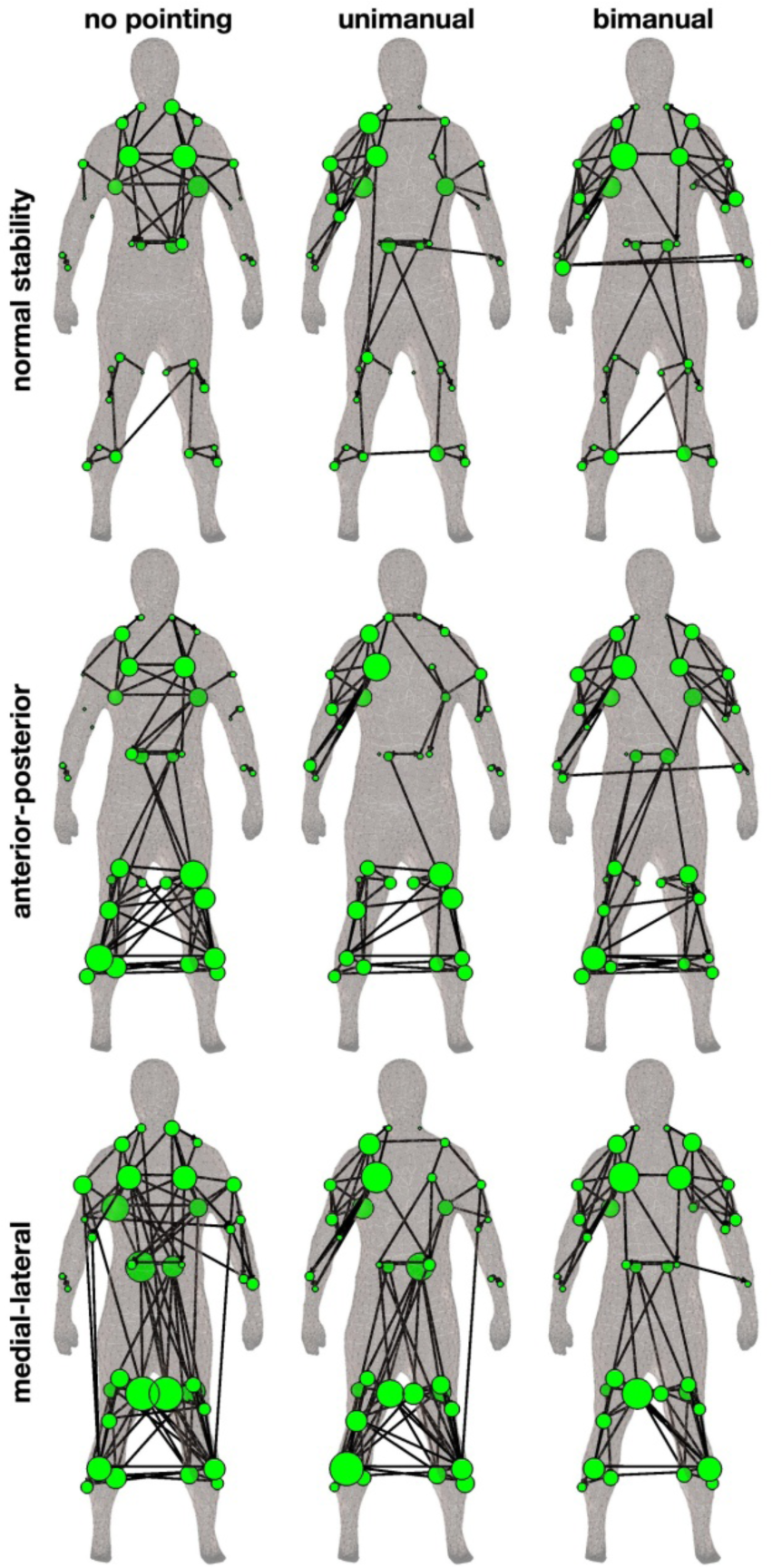
Directed muscle networks across conditions estimated using conditional transfer entropy. Arrows indicate significant interactions and node size represents the in-degree of the significant edges. The in-degree is estimated in group-level networks.

To test whether these changes were statistically significant, we compared the in-degree of each node across conditions using repeated-measures ANOVA with Bonferroni-Holm correction for multiple comparisons. We found a significant main effect of pointing behavior in three muscles in the upper body (right and left PMA and left LD) and a significant main effect of postural instability in thirteen muscles mainly located in the lower body (right and left TA, SOL, RF, VL, AL, the left BF and the right EO and LO, Fig. 5A). We selected the muscles showing the largest main effects to investigate the experimental effects in more detail. The PMA situated at the chest of the body revealed the largest effect of pointing behavior on the in-degree of transfer entropy (right: F_(2,117)_=18.8, P_corr_<0.0001; left: F_(2,117)_=10.8, P_corr_=0.005). Conditional transfer entropy was largest in the no pointing condition and reduced in unimanual and bimanual pointing conditions (Fig. 5B). This pattern was also mirrored in the total transfer, i.e. the global directed effects from all muscles towards the target. Information storage, i.e. the presence of predictable/repeatable temporal patterns in the EMG envelopes, did not show a clear effect of pointing condition. Finally, total information, which is closely related to the variance of the EMG envelopes, was strongly enhanced when the corresponding arm was involved in the pointing task, i.e. during unimanual and bimanual pointing for the right PMA and only during bimanual pointing for the left PMA. For postural instability, we observed the largest effect in the AL muscle located on the medial side of the thigh (right: F_(2,117)_=61.9, P_corr_<0.0001; left: F_(2,117)_=27.0, P_corr_<0.0001). Conditional transfer entropy and total transfer were lowest in the normal standing condition, increased during anterior-posterior instability and highest during medial-lateral instability (Fig. 5B). A similar pattern was observed for total storage and information. These information metrics hence showed distinct effects in these two muscles across experimental conditions.

**Figure 5.**
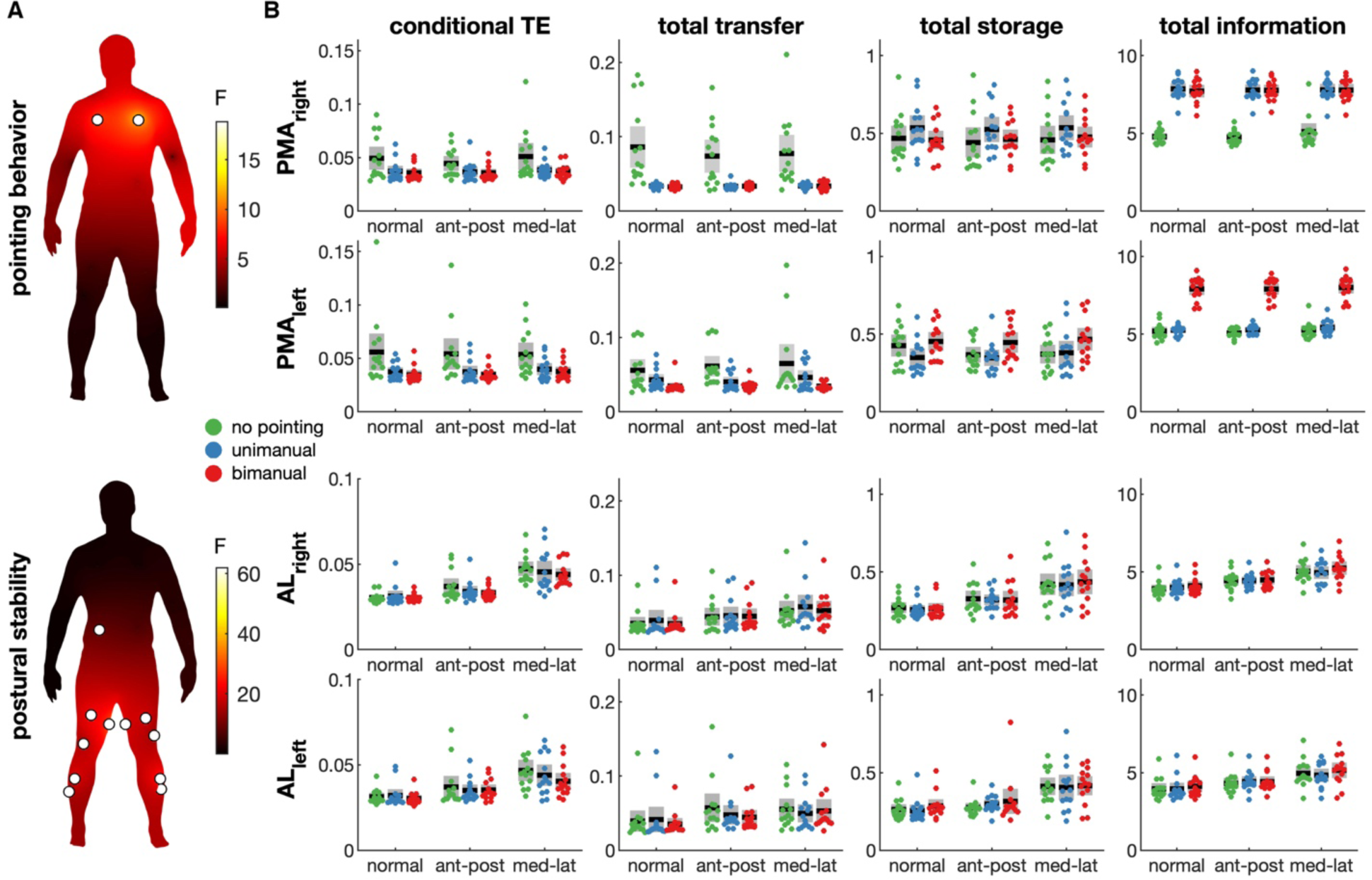
Comparison of information metrics across experimental conditions. **A)** Statistical results of two-way ANOVA of the weighted in-degree of conditional transfer entropy. The F-statistic is depicted as color-coded values at the three-dimensional coordinates of the locations where the muscle activity was recorded and interpolated on the other points of the body surface mesh (Makarov et al., 2015). The top panel shows the main effect of pointing behavior and the bottom panel the main effect of postural stability. White circles indicate nodes (muscles) with a significant effect after Bonferroni-Holm correction. **B)** Descriptive statistics of three information metrics (conditional transfer entropy, total information transfer and total information) for selected muscles (PMA left, right and AL left, right). Dots show values of individual participants, the black line the group averages, and the grey boxes the standard error.

## 4. Discussion

We used information decomposition of multichannel EMG to investigate task-related changes in the functional coordination of muscles distributed across the body. Participants performed different postural tasks in which pointing behavior (no, unimanual and bimanual pointing) and postural stability (normal, anterior-posterior and medial-lateral instability) were experimentally manipulated. Undirected and directed muscle networks were assessed by estimating mutual information and information transfer between EMG envelopes. A sparse network was observed when defining the edges based on statistically significant interactions, in particular when conditioning for information from other muscles. We compared muscle networks across conditions using the weighted in-degree of conditional transfer entropy and observed that task-related effects were confined to the muscles that are involved in the task: pointing behavior affected upper body muscles and postural behavior mainly affected lower body muscles. Furthermore, information decomposition revealed distinct patterns underlying both effects. Manual and bimanual pointing were associated with reduced transfer to the PMA muscles, but an increase in total information compared to no pointing, while postural instability resulted in increased information transfer, storage and information in the AL muscles compared to normal stability. These findings show that information decomposition and network analysis of surface EMG can be used as a tool to map changes in functional interactions in the sensorimotor system during postural control tasks.

We used predictive information measures to study the neural circuitry interconnecting motor neuron pools using the EMG activity of several muscles distributed across the body. Such measures allow quantifying how much of the uncertainty about the current state of a muscle is reduced by the knowledge of the past states visited by the whole muscle network, and to distinguish between the sources of the predictive information relevant to a target muscle: whether it originates from the past of the target, from a source muscle, or from other muscles within the network (Fig. 1D). The decomposition of predictive information into information storage and transfer, and the distinction between bivariate and conditional information transfer, provide a more detailed picture of the activity of each node and the connectivity between nodes in a dynamic network (Lizier, 2012). This has been shown in brain networks (Wibral et al., 2015), physiological networks (Faes et al, 2017a), and networks of brain-body interactions (Zanetti et al., 2019), and here it is documented for the first time in muscle networks estimated from surface EMG. In particular, we assessed conditional mutual information and transfer entropy to account for other variables that may influence the relationship between source and target. Differentiation between spurious, indirect, and causal associations in observational data has been a long-standing problem and requires to control for confounding variables (Greenland et al., 1999; Grimes and Schulz, 2002). Here, we used a fully multivariate approach where we examined the potential effect of one variable on another while simultaneously controlling for the effect of many other variables. The use of a linear VAR models puts the computed indexes in the well-framed context of Granger causality (Seth et al., 2015) and greatly enhances computational reliability (Barnett and Seth, 2015). We observed a sparser network when estimating edges weights using conditional information metrics, which shows that the conventional unconditioned metrics indeed conflate direct and indirect associations between muscles, as well as the effects of common drivers.

Although we performed multivariate information decomposition of 36 EMG signals and thus accounted for a large number of confounding variables, a confounding bias may also arise from exogenous (unrecorded) variables. A key candidate would be supraspinal motor areas such as the motor cortex that has direct and indirect projections to the spinal motor neurons via descending pathways (Lemon, 2008). It has been well-established that activity from sensorimotor cortex and muscle activity are correlated, in particular in the beta band (15-30 Hz) (Conway et al., 1995; Mima and Hallett, 1999) but also at low frequencies (< 3 Hz) (Bourguignon et al., 2017), and that corticospinal drive results in correlated activity of spinal motor neurons innervating functionally related muscles (Farmer et al., 1993). Hence, an obvious extension of the current approach would be to simultaneously record multichannel EMG and EEG and apply multivariate information decomposition to the combined dataset to distinguish between intermuscular and corticomuscular associations (Boonstra, 2013). Directed connectivity has been used to assess unidirectional connectivity between motor cortex and leg muscles during walking (Artoni et al., 2017). Multivariate information decomposition of combined EEG and EMG data would be a principled approach to study the brain-muscle networks involved in human motor coordination.

Another possible limitation is that we assessed functional interactions in the time domain. Time domain analysis is widely used to study motor coordination, for example by extracting muscle synergies from the EMG envelopes (d’Avella et al., 2003; Tresch et al., 2006), but spectral analysis has shown that rhythmic activity in the human motor system can be observed at multiple distinct frequencies (McAuley and Marsden, 2000). We previously showed that intermuscular coherence between postural muscles at different frequencies revealed distinct network topologies, which indicate the functioning of a multiplex network organization (Kerkman et al., 2018). Multivariate frequency-domain analysis of coupled processes can be used to assess frequency-dependent causal interactions in physiological time series (Faes and Nollo, 2011) and we have applied this to assess frequency-dependent directed muscle networks (Boonstra et al., 2015). Nevertheless, network information measures have the advantage that they can be conveyed in a framework that identifies the basic components of information processing (i.e., storage, transfer and modification) into which network dynamics are dissected (Lizier, 2012), a perspective which is not available in the frequency domain. Moreover, information measures provide a compact description of the dynamics which subsumes oscillatory behaviors in different frequency bands. For instance, an increased information storage, such as observed in the AL muscles while perturbing postural stability, likely reflects the emergence of dominant oscillatory activity in specific frequency bands, which is also related to higher intermuscular coherence (Kerkman et al., 2018). Finally, electrical cross-talk may have affected the information metrics estimated between EMGs (Farina et al., 2014), in particular as functional connectivity was mainly observed between neighboring muscles. However, in a previous study using the same data set (Kerkman et al., 2018), we did not find elevated coherence levels across frequencies indicative of cross-talk. As cross-talk results in instantaneous correlations between signals, the effect on time-lagged interactions will be reduced. In addition, we found interactions between bilateral muscles in the arms and legs, consistent with previous studies (Boonstra et al., 2015; de Vries et al., 2016), which cannot be caused by cross-talk. We therefore expect the effects of cross-talk to be minor.

The current findings show that directed interactions between muscles are widespread and task-dependent. There is extensive evidence that motor neuron pools innervating muscles that are anatomically or functionally related receive correlated inputs (Bremner et al., 1991; Farmer et al., 1993; Gibbs et al., 1995; Laine et al., 2015). These correlated inputs are generally thought to originate from supraspinal areas through divergent pathways or from presynaptic synchronization, i.e. synchronization of the neuronal populations that project to the spinal motor neuron pool (Kirkwood, 2016). The results showing clear patterns of conditional transfer entropy between EMGs suggest the involvement of additional pathways that interconnect different motor neuron pools. A spinal network of interneurons interconnecting motor neuron pools within the spinal cord is involved in the coordination of muscle activity (Levine et al., 2014; Takei et al., 2017). Moreover, reflex pathways are a well-known mechanism whereby muscle activity from different muscles influences each other (Latash, 2007). While reflexes are largely automated, modern views suggest that gain modulation of reflex pathways are also involved in the control of voluntary movement (Azim and Seki, 2019; Krakauer, 2019). In particular, it is now thought that spinal interneurons control the input that spinal motor neurons receive from different primary afferents and descending tracts and that the brain modulates the activity of these spinal interneurons to control movement (Pierrot-Deseilligny and Burke, 2012). According to the equilibrium point hypothesis, the central nervous system indeed controls movements by setting the threshold or gain of these spinal reflexes (Latash, 2008). An interesting hypothesis that can be tested in future studies is that the task-related changes in functional interactions between muscles observed in this study reflects the change in gain of spinal reflex pathways.

## Acknowledgements

TB was supported by a Future Fellowship from the Australian Research Council (FT180100622). JK was supported by the Netherlands Organization for Scientific Research (NWO 016.156.346 awarded to Nadia Dominici) and thanks Nadia Dominci and Andreas Daffertshofer for their supervision. The computational resources (Stevin Supercomputer Infrastructure) and services used in this work were provided by the VSC (Flemish Supercomputer Center), funded by Ghent University, FWO and the Flemish Government – department EWI. We thank Frederik Van de Steen for helping to set up the analyses on said infrastructure.

